# Delta spike P681R mutation enhances SARS-CoV-2 fitness over Alpha variant

**DOI:** 10.1101/2021.08.12.456173

**Authors:** Yang Liu, Jianying Liu, Bryan A. Johnson, Hongjie Xia, Zhiqiang Ku, Craig Schindewolf, Steven G. Widen, Zhiqiang An, Scott C. Weaver, Vineet D. Menachery, Xuping Xie, Pei-Yong Shi

## Abstract

SARS-CoV-2 Delta variant has rapidly replaced the Alpha variant around the world. The mechanism that drives this global replacement has not been defined. Here we report that Delta spike mutation P681R plays a key role in the Alpha-to-Delta variant replacement. In a replication competition assay, Delta SARS-CoV-2 efficiently outcompeted the Alpha variant in human lung epithelial cells and primary human airway tissues. Delta SARS-CoV-2 bearing the Alpha-spike glycoprotein replicated less efficiently than the wild-type Delta variant, suggesting the importance of Delta spike in enhancing viral replication. The Delta spike has accumulated mutation P681R located at a furin cleavage site that separates the spike 1 (S1) and S2 subunits. Reverting the P681R mutation to wild-type P681 significantly reduced the replication of Delta variant, to a level lower than the Alpha variant. Mechanistically, the Delta P681R mutation enhanced the cleavage of the full-length spike to S1 and S2, leading to increased infection via cell surface entry. In contrast, the Alpha spike also has a mutation at the same amino acid (P681H), but the spike cleavage from purified Alpha virions was reduced compared to the Delta spike. Collectively, our results indicate P681R as a key mutation in enhancing Delta variant replication via increased S1/S2 cleavage. Spike mutations that potentially affect furin cleavage efficiency must be closely monitored for future variant surveillance.

## Introduction

The continuous emergence of new variants of severe acute respiratory syndrome coronavirus 2 (SARS-CoV-2) poses the greatest threat to pandemic control, vaccine effectiveness, therapeutic efficacy, and surveillance. Since its emergence in late 2019, mutations have unceasingly emerged in the circulating viruses, leading to variants with enhanced transmissibility, evasion of therapeutic antibodies, and breakthrough infections in vaccinated individuals.^1–6^ Since the viral spike glycoprotein is responsible for binding to the human cellular receptor angiotensin-converting enzyme (ACE2), many mutations have accumulated in the spike gene with the potential to alter viral fitness or to escape immunity. The variants have emerged from different geographic regions and, depending on their biological properties, spread to other regions. The World Health Organization (WHO) has classified variants as “variants of concern” *(i.e.*, Alpha, Beta, Gamma, and Delta) and “variants of interest” *(i.e.*, Eta, Iota, Kappa, and Lambda).^7^ The Alpha variant was first identified in the United Kingdom in September 2020 and subsequently became dominant in many parts of the world. Afterwards, the Delta variant emerged in India in October 2020 and has now spread to over 119 countries, displacing the Alpha variant globally.^7,8^ From May 2 to July 31 of 2021, the prevalence of the Delta variant in the USA had increased from 1.3% to 94.4%, whereas the prevalence of the Alpha variant had decreased from 70% to 2.4%. More seriously, the Delta variant has been associated with increased transmissibility, disease severity, and breakthrough infections in vaccinated individuals.^5,9–11^ The mutation(s) that have driven the explosive spread of the Delta variant and its displacement of the Alpha variant remain to be defined. In this study, we used a reverse genetic approach to identify the molecular determinant(s) for the enhanced fitness of Delta variant and its dominance over the Alpha variant.

## Results

We constructed infectious cDNA clones for the Alpha (GISAIS ID: EPI_ISL_999340) and Delta (GISAIS ID: EPI_ISL_2100646) SARS-CoV-2 variants using a previously established protocol (**Extended data Fig. 1**).^12,13^ The infectious cDNA clones enabled us to prepare recombinant Alpha and Delta SARS-CoV-2 variants (**Fig. 1a**). Both Alpha and Delta variants rescued from these clones developed smaller plaques on Vero E6 cells than the earlier USA/WA1-2020 (wild-type) strain isolated in January 2020 (**Extended data Fig. 2**). Sequencing analysis showed no undesired mutations in the rescued recombinant virus stocks. To compare the viral replication fitness between the Alpha and Delta variants, we performed a competition assay by infecting cells with a mixture of the two viruses at a plaque-forming unit (PFU) ratio of 1:1, followed by quantifying the ratios of the two viral RNA species at different days post infection. Compared with analyzing individual viruses separately, the competition assay has the advantages of (i) a built-in internal control of each viral replication and (ii) elimination of host-to-host variation that reduces experimental power. Due to its precision and reproducibility^14^, the competition assay has been widely used to study microbial fitness,^15–17^ including SARS-CoV-2^1,18^. When infecting human lung adenocarcinoma Calu-3 cells, the RNA ratio of Delta versus Alpha increased to 3.0, 7.0, and 4.1 at 24, 36, and 48 h post infection, respectively (**Extended data Fig. 3**). When infecting primary human airway epithelial (HAE) cultures, the RNA ratio of Delta versus Alpha increased from 1.7 on day 1 to 3.1 on day 5 (**Fig. 1b**). These results indicate that Delta variant has greater replication fitness compared to the Alpha variant in *in vitro* respiratory models of SARS-CoV-2 infection.

**Figure 1.**
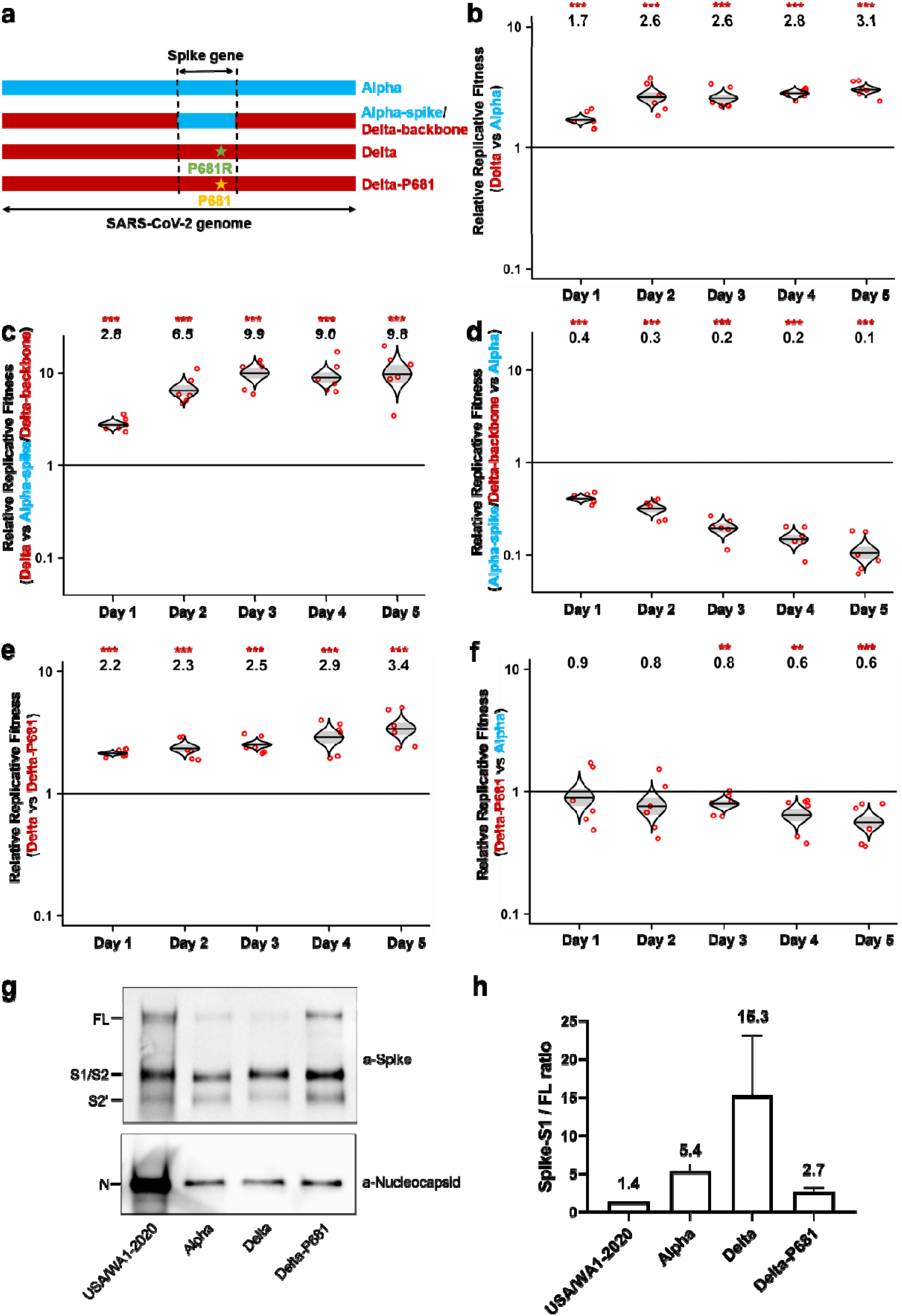
Delta P681R mutation enhances SARS-CoV-2 fitness over Alpha variant through improving spike protein processing. **a,** Schemes of Alpha variant, Delta variant, Delta variant bearing Alpha-spike, and Delta variant with a P681R-to-P681 reversion. The spike gene of Delta variant was swapped with Alpha variant, resulting in chimeric SARS-CoV-2 “Alpha-spike/Delta-backbone.” The Delta P681R mutation was reverted wild-type P681, resulting in “Delta-P681” virus. Blue and red colors indicate Alpha and Delta variants, respectively. **b-f,** Viral replication competitions among Alpha, Delta, Alpha-spike/Delta-backbone, and Delta-P681 viruses on primary human airway epithelial (HAE) cells. Equal PFU of two viruses were mixed and inoculated onto HAE cells at an MOI of 5. Five pairs of viral competition are presented: Delta and Alpha (**b**), Delta and Alpha-spike/Delta-backbone (**c**), Alpha-spike/Delta-backbone and Alpha (**d**), Delta variant and Delta-P618 (**e**), and Delta-P681 and Alpha (**f**). After 2 h incubation, the cells were washed thrice with DPBS and maintained for 5 days. The secreted viruses were collected daily in DPBS after incubation at 37°C for 30 min. Red dots represent individual cell cultures (n=6), the horizontal lines in each catseye represent the mean, shaded regions represent standard error of the mean; y-axes use a log_10_ scale. Black numbers above each set of values (catseye) indicate the ratios of two viral RNA species. *P* values are calculated for group coefficient using linear regression model. \*\**p*<0.01, \*\*\**p*<0.001. **g,** Spike cleavages of purified virions. USA/WA1-2020, Alpha, Delta, and Delta-P681 viruses were purified and analyzed by Western blot using polyclonal antibodies against spike and anti-nucleocapsid antibodies. Full-length spike (FL), cleaved S1/S2, and S2’ proteins were annotated. One representative image of two experiments is shown. **h,** Quantification of spike processing of different variants from **g**. Densitometry was performed to quantify the cleavage efficiency of FL spike to S1/S2 subunits using ImageLab 6.0.1. The ratios of S1/S2 over FL were calculated to indicate spike processing efficiencies. The average results of two experiments were presented as mean ± standard deviation.

To examine if the spike gene alone determines the improved replication fitness of the Delta variant, we constructed a chimeric Delta SARS-CoV-2 bearing the Alpha-spike glycoprotein (*i.e.*, Alpha-spike/Delta-backbone virus; **Fig. 1a** and **Extended data Fig. 2b**). In a competition assay on HAE culture, the RNA ratio of Delta versus Alpha-spike/Delta-backbone continuously increased from 2.8 on day 1 to 9.8 on day 5 post infection (**Fig. 1c**), suggesting that (i) the Alpha-spike reduces the replication fitness of the Delta variant and (ii) the spike gene drives the improved replication of Delta variant. Interestingly, the Alpha-spike/Delta-backbone virus replicated less efficiently than the Alpha variant on HAE culture (**Fig. 1d**), suggesting that, in contrast to Delta spike mutations that enhance replication, mutations outside the spike gene of the Delta variant reduced to some degree fitness for viral replication. The above quantifications of viral competition outcomes were measured by Sanger sequencing (**Fig. 1b-d**) and subsequently verified by Illumina next generation sequencing (NGS; **Extended data Fig. 4**).

Delta spike has accumulated mutations T19R, G142D, E156G, F157-R158 deletion, L452R, T478K, D614G, P681R, and D950N,^19^ among which P681R is located at a furin cleavage site (<u>P</u>RRAR↓S with P681 underlined and “↓” indicating furin cleavage) that is absent in other group 2B coronaviruses.^20^ Since the furin cleavage site was shown to be important for SARS-CoV-2 replication and pathogenesis,^21,22^ we hypothesized that mutation P681R may improve the furin cleavage efficiency of full-length spike to S1 and S2, leading to a more efficient virus entry into respiratory epithelial cells. To test this hypothesis, we reverted the Delta P681R mutation to wild-type P681 in the Delta SARS-CoV-2 (**Extended data Fig. 2a**). The Delta-P681 virus developed equivalent plaque morphology compared with Delta (**Extended data Fig. 2b**). Remarkably, the P681 reversion attenuated Delta variant replication on HAE cultures, as evidenced by the increase in the RNA ratio of wild-type Delta versus Delta-P681 from 2.2 on day 1 to 3.4 on day 5 (**Fig. 1e**). The replication of the Delta-P681 virus was even lower than that of the Alpha variant, as suggested by the decrease in the RNA ratio of Delta-P681 versus Alpha variant from 0.9 on day 1 to 0.6 on day 5 (**Fig. 1f**). These results demonstrate that mutation P681R at the furin cleavage site plays a critical role in enhancing the replication of the Delta variant on primary human airway cultures.

We directly evaluated the spike cleavage of Alpha, Delta, Delta-P681, and wild-type SARS-CoV-2. Virions were prepared from Vero E6 cells expressing TMPRSS2, a host serine protease that is required for SARS-CoV-2 entry via the ACE2-mediated cell surface mechanism.^23^ After virions were purified through sucrose cushion ultracentrifugation, pelleted viruses were analyzed for spike cleavage by Western blotting (**Fig. 1g**). The results showed that spike processing efficiency of the purified virions occurred in the order of Delta > Alpha > Delta-P681 > wild-type virions, with ratios of S1 versus full-length spike of 15.3, 5.4, 2.7, and 1.4, respectively (**Fig. 1h**). It should be noted that the Alpha variant also has a spike mutation at amino acid position 681 (P681H), which may contribute to the increase in spike cleavage when compared with the wild-type USA/WA1-2020 virus; however, a recent study showed that mutation P681H alone did not enhance viral fitness or transmission.^4^ Overall, our results indicate a correlation of improved spike cleavage with enhanced viral replication of Delta variant.

To exclude the possibility that enhanced replication of the Delta variant over Alpha was due to an improved spike/ACE2 receptor interaction, we performed a binding assay using recombinant spike receptor-binding domain (RBD) and human ACE2 proteins on a Bio-Layer Interferometry (BLI) system (**Extended data Fig. 5**). Within the RBD, the Alpha RBD has a N501Y mutation, whereas Delta RBD has L452R and T478K mutations. The BLI results indicate that the Alpha RBD has a higher binding affinity for ACE2 than Delta RBD, as indicated by >200-fold K_D_ improvement (**Extended data Fig. 5**). These data strongly argue that the higher replication of Delta variant than Alpha variant is not due to an improved spike/ACE2 receptor binding.

## Discussion

Since the emergence of SARS-CoV-2, the virus has accumulated mutations improving fitness and transmission. First, it accumulated a D614G mutation in the spike gene to enhance viral transmission.^1,3,6,24^ This mutation promotes spike RBD in an “open” conformation to facilitate ACE2 receptor binding.^25^ Subsequently, another spike mutation N501Y emerged independently in Alpha, Beta, and Gamma variants from the United Kingdom, South Africa, and Brazil, respectively. The N501Y mutation further increases the binding affinity between the spike protein and ACE2, leading to additional improvement in viral transmission.^4,10,26^ Most recently, the Delta variant emerged and spread explosively, replacing the Alpha variant around the world. The current study demonstrates that spike mutation(s) are responsible for the enhanced viral fitness of the Delta variant over Alpha. Importantly, the unique P681R mutation plays a critical role in this fitness advantage and increases the processing of Delta spike to S1 and S2, most likely through an improved furin cleavage when newly assembled virions egress through the trans-Golgi network. Although the original SARS-CoV-2 strain has a functional furin cleavage site with a minimal recognition site of RXXR↓,^27^ adjacent residues influence the cleavage efficiency.^28^ The P681R substitution clearly augments spike processing and is likely the main driver of the fitness advantage observed in Delta variant. When the Delta variant infects respiratory epithelial cells, it binds to ACE2 receptor via the RBD in S1. Already cleaved at the S1/S2 site, the Delta virion facilitates cleavage at S2’ by the cell surface protease TMPRSS2, leading to an activation of the S2 fusion peptide (FP) for viral and plasma membrane fusion.^23^ Thus, the improved spike cleavage enhances viral replication when the Delta variant infects respiratory epithelial cells.

One major concern with the emergence of Delta variant is its association with increased breakthrough infections in vaccinated people.^5,9^ As a critical target for host immunity, changes in the spike protein of SARS-CoV-2 variants have been implicated in reducing antibody neutralization.^29^ Among all tested variants (including Delta), the Beta and Kappa variants have been shown to reduce the BNT162b2 vaccine-elicited neutralizing titers the most;^2,30–32^ yet, BNT162b2 showed 100% vaccine efficacy against Beta variant-associated severe, critical or fatal disease in Qatar and 100% real-world effectiveness against Beta variant-associated COVID-19 in South Africa.^33,34^ These real-world vaccine efficacy/effectiveness and *in vitro* neutralization results argue that breakthrough infections by the Delta variant in vaccinated individuals do not reflect immune escape. Instead, the increased breakthrough infection is likely due to enhanced viral replication fitness of the Delta variant through augmented spike processing. Consistent with this hypothesis, the viral RNA loads in the oropharynx from Delta variant-infected patients were >1,200-fold higher than those from the original Wuhan virus infected patients.^9^ Together, the results indicate changes in viral processing and infection efficiency, rather than evasion of antibodies, drive breakthrough infections of Delta variant in vaccinated individuals.

In summary, using a reverse genetic system and primary human airway cultures, we have identified spike mutation P681R as a significant determinant for enhanced viral replication fitness of the Delta compared to the Alpha variant. The P681R mutation enhances spike protein processing through the improved furin cleavage site. As new variants continue to emerge, spike mutations that affect furin cleavage efficiency, as well as other mutations that may increase viral replication, pathogenesis, and/or immune escape, must be closely monitored.

## Methods

### Cells

African green monkey kidney epithelial Vero E6 cells (ATCC, Manassas, VA, USA) were grown in Dulbecco’s modified Eagle’s medium (DMEM; Gibco/Thermo Fisher, Waltham, MA, USA) with 10% fetal bovine serum (FBS; HyClone Laboratories, South Logan, UT) and 1% penicillin/streptomycin (Gibco). Human lung adenocarcinoma epithelial Calu-3 cells (ATCC) were maintained in a high-glucose DMEM containing sodium pyruvate and GlutaMAX (Gibco) with 10% FBS and 1% penicillin/streptomycin at 37°C with 5% CO_2_. The EpiAirway system is a primary human airway 3D tissue model purchased from MatTek Life Science (Ashland, MA, USA). This EpiAirway system was maintained with the provided culture medium at 37°C with 5% CO_2_ following manufacturer’s instruction. All other culture medium and supplements were purchased from Thermo Fisher Scientific (Waltham, MA, USA). All cell lines were verified and tested negative for mycoplasma.

### Construction of infectious cDNA clones and SARS-CoV-2 mutant viruses

The full-length (FL) cDNA clones of Alpha and Delta variants were constructed through mutagenesis of a previously established cDNA clone of USA/WA1-2020 SARS-CoV-2.^12,13^ The previous seven-fragment *in vitro* ligation method was improved to a three-fragment ligation approach (**Extended data Fig. 1a**) to construct the full-length cDNA clones of Alpha and Delta SARS-CoV-2, resulting in Alpha-FL and Delta-FL, respectively. Prior to the three-fragment ligation, mutations from Alpha or Delta variants were engineered into individual fragments of USA/WA1-2020 using a standard mutagenesis method. The sequences for constructing Alpha, Delta and Alpha-spike/Delta-backbone were downloaded from GISAID database, the accession ID for Alpha is EPI_ISL_999340, accession ID for Delta is EPI_ISL_2100646. Individual point mutations for Alpha (NSP3: P153L, T183I, A890D, I1412T; NSP6: SGF106-108del; NSP12: P323L; Spike: HV69-70del, Y145del, N501Y, A570D, D614G, P681H, T716I, S982A, D1118H; ORF8: Q27stop, R52I, Y73C, S84L; N: D3L, R203K, G204R, S235F) and individual point mutations for Delta (NSP2: P129L; NSP3: P822L; H1274Y; NSP4: A446V; NSP6: V149A; NSP12: P323L; V355A; G671S; NSP13: P77L; NSP15: H234Y; Spike: T19R, G142D, E156G, FR157-158del, L452R, T478K, D614G, P681R, D950N; ORF3a: S26L; M: I82T; ORF7a: V82A; L116F; T120I; ORF8: S84L; DF119-120del; N: D63G; R203M; D377Y; R385K) were introduced into subclones of individual fragments by overlapping fusion PCR. For preparing Alpha-spike/Delta-backbone virus, the spike gene of Delta was replaced with the spike gene of the Alpha. For preparing Delta-P681 virus, the P681 reversion was introduced into a subclone containing Delta spike gene by overlapping fusion PCR. All primers used for the construction were listed in **Extended Data Table 1**. The full-length infectious clones of SARS-CoV-2 variants were assembled by *in vitro* ligation of contiguous DNA fragments. *In vitro* transcription was then performed to synthesize full-length genomic RNA. For recovering recombinant viruses, the RNA transcripts were electroporated into Vero E6 cells. The viruses from electroporated cells were harvested at 40 h post electroporation and served as P0 stocks. All viruses were passaged once on Vero E6 cells to produce P1 stocks for subsequent experiments. All P1 viruses were subjected to next generation sequencing to confirm the introduced mutations without undesired changes. Viral titers were determined by plaque assay on Vero E6 cells. All virus preparation and experiments were performed in a biosafety level 3 (BSL-3) facility. Viruses and plasmids are available from the World Reference Center for Emerging Viruses and Arboviruses (WRCEVA) at the University of Texas Medical Branch.

### RNA extraction, RT-PCR, and Sanger sequencing

Cell culture supernatants were mixed with a five-fold excess of TRIzol™ LS Reagent (Thermo Fisher Scientific, Waltham, MA, USA). Viral RNAs were extracted according to the manufacturer’s instructions. The RNAs were amplified using a SuperScript™ III One-Step RT-PCR kit (Invitrogen, Carlsbad, CA, USA) following the manufacturer’s protocol. The size of desired amplicon was verified with 2 μl of PCR product on an agarose gel. The remaining 18 μl of RT-PCR DNA was purified by a QIAquick PCR Purification kit (Qiagen, Germantown, MD, USA). Sequences of the purified RT-PCR products were generated using a BigDye Terminator v3.1 cycle sequencing kit (Applied Biosystems, Austin, TX, USA). The sequencing reactions were purified using a 96-well plate format (EdgeBio, San Jose, CA, USA) and analyzed on a 3500 Genetic Analyzer (Applied Biosystems, Foster City, CA). The peak electropherogram height representing each mutation site and the proportion of each competitor was analyzed using the QSVanalyser program. For the competition assay, R software is used for the figure generation and statistical analysis. The presented viral RNA ratios in the figures were normalized by the input viral RNA ratios (**Extended Data Table 2**).

### Plaque assay

Approximately 1.2×10^6^ Vero E6 cells were seeded to each well of 6-well plates and cultured at 37°C, 5% CO_2_ for 16 h. Virus was serially diluted in DMEM with 2% FBS and 200 μl diluted viruses were transferred onto the cell monolayers. The viruses were incubated with the cells at 37°C with 5% CO_2_ for 1 h. After the incubation, overlay medium was added to the infected cells per well. The overlay medium contained DMEM with 2% FBS, 1% penicillin/streptomycin, and 1% sea-plaque agarose (Lonza, Walkersville, MD). After 2.5-day incubation, plates were stained with neutral red (Sigma-Aldrich, St. Louis, MO, USA) and plaques were counted on a light box.

### Next generation sequencing (NGS)

The competition results generated by Sanger sequencing were confirmed using NGS methods. Briefly, viral RNA samples from competition groups of (i) Delta versus Alpha and (ii) Delta versus Alpha-spike/Delta-backbone were used for a specific one-step RT-PCR that containing the A23063T mutation site. Viral RNA samples from competition group of Alpha versus Alpha-spike/Delta-backbone were quantified by the T14444C mutation. The RT-PCR primers were listed in **Extended Data Table 1**. The PCR products were purified by a QIAquick PCR Purification kit (Qiagen, Germantown, MD) according to the manufacturer’s protocol. Dual-indexed adapter sequences (New England BioLabs, Ipswich, MA) were added with 5 cycles of PCR. Samples were pooled and sequenced on an Illumina MiniSeq Mid-Output flow cell with the paired-end 150 base protocol. The reads were filtered for Q-scores of 37 at the A23063T and T14444C mutation sites and adjacent bases and counted. For the competition assay, R software is used for the figure generation and statistical analysis.

### Viral infection of cell lines

Approximately 3×10^5^ Calu-3 cells were seeded onto each well of 12-well plates and cultured at 37°C, 5% CO_2_ for 16 h. Equal PFUs of two viruses were inoculated onto Calu-3 cells at a final MOI of 0.1. The mixed viruses were incubated with the cells at 37°C for 2 h. After infection, the cells were washed thrice with DPBS to remove residual viruses. One milliliter of culture medium was added into each well. At each time point, 100 μl of culture supernatants were lysed in TRIzol LS reagent for the detection of competition assay, and 100 μl of fresh medium was added into each well to replenish the culture volume. The cells were infected in triplicate for each group of viruses. All samples were stored at −80°C until analysis.

### Primary human airway cultures

The EpiAirway system is a primary human airway 3D mucociliary tissue model consisting of normal, human-derived tracheal/bronchial epithelial cells. Different combinations of mixed viruses for competition assays were inoculated onto the culture at a total MOI of 5. After 2 h infection at 37 °C with 5% CO_2_, the inoculum was removed, and the culture was washed three times with DPBS. The infected epithelial cells were maintained without any medium in the apical well, and medium was provided to the culture through the basal well. The infected cells were incubated at 37 °C, 5% CO_2_. From day 1 to day 5 post infection, 300 μl of DPBS were added onto the apical side of the airway culture and incubated at 37°C for 30 min to elute progeny viruses. All virus samples in DPBS were stored at −80°C and quantified by plaque assays on Vero E6 cells.

### Virion purification and Western blotting

Vero E6 expressing TMPRSS2 were infected with different SARS-CoV-2 at an MOI of 0.01. At 24 h post infection, the culture medium was collected, purified through a 20% sucrose cushion, and analyzed by Western blot as previously described.^21^ Densitometry was performed to quantify the cleavage efficiency of full-length spike to S1/S2 subunits using ImageLab 6.0.1 (Bio-Rad #12012931). The average results of two experiments were presented.

### Spike RBD and ACE2 binding

The human ACE2 protein was purchased from Sino Biological (Beijing, China; Cat# 10108-H08H) and the human IgG1 Fc-tagged RBD proteins were made in-house using a method as previously described^35^. The affinity measurement was performed on the ForteBio Octet RED 96 system (Sartorius, Goettingen, Germany). Briefly, the RBD proteins (20 μg/ml) of Alpha or Delta RBDs were captured onto protein A biosensors for 300s. The loaded biosensors were then dipped into the kinetics buffer for 10 s for adjustment of baselines. Subsequently, the biosensors were dipped into serially diluted (from 1.23 to 300 nM) human ACE2 protein for 200 s to record association kinetics and then dipped into kinetics buffer for 400 s to record dissociation kinetics. Kinetic buffer without ACE2 was used to correct the background. The Octet Data Acquisition 9.0 software was used to collect affinity data. For fitting of K_D_ values, Octet Data Analysis software V11.1 was used to fit the curve by a 1:1 binding model using the global fitting method.

### Statistics

For virus competition experiments, relative replicative fitness values for different variants were analyzed according to w=(f0/i0), where i0 is the initial two-virus ratio and f0 is the final two-virus ratio after competition. Sanger sequencing (initial timepoint T0) counts for each virus being compared were based upon average counts over three replicate samples of inocula per experiment, and post-infection (timepoint T1) counts were taken from samples of individual subjects. Multiple experiments were performed, so that f0/i0 was clustered by experiment. To model f0/i0, the ratio T0/T1 was found separately for each subject in each virus group, log (base-10) transformed to an improved approximation of normality and modeled by analysis of variance with relation to group, adjusting by experiment when appropriate to control for clustering within experiment. Specifically, the model was of the form Log10_CountT1overCountT0 ∼ Experiment + Group. Fitness ratios between the two groups [the model’s estimate of w=(f0/i0)] were assessed per the coefficient of the model’s Group term, which was transformed to the original scale as 10^coefficient. This modeling approach compensates for any correlation due to clustering within experiment similarly to that of corresponding mixed effect models and is effective since the number of experiments was small. Statistical analyses were performed using R statistical software (R Core Team, 2019, version 3.6.1). In all statistical tests, two-sided alpha=.05. Catseye plots^36^, which illustrate the normal distribution of the model-adjusted means, were produced using the “catseyes” package^37^.

## Data availability

Extended Data and source data for generating main figures are available in the online version of the paper. Any other information is available upon request.

## Acknowledgments

P.-Y.S. was supported by National Institutes of Health (NIH) grants AI134907 and UL1TR001439, and awards from the Sealy and Smith Foundation, the Kleberg Foundation, the John S. Dunn Foundation, the Amon G. Carter Foundation, the Gillson Longenbaugh Foundation, and the Summerfield Robert Foundation. Z.A. was supported in by a Welch Foundation grant AU-0042-20030616 and Cancer Prevention and Research Institute of Texas (CPRIT) Grants RP150551 and RP190561. S.C.W. was supported by NIH grant R24 AI120942. V.D.M. was supported by NIH grants AI153602 and 1R21AI145400. J.L. and B.A.J. were supported by James W. McLaughlin Fellowship Fund.

## Author contributions

Conceptualization, Y.L., J.L., S.C.W., V.D.M., X.X., P.-Y.S.; Methodology, Y.L., J.L., B.A.J., H.X., Z.K., C.S., S.G.W., Z.A., X.X.; Investigation, Y.L., J.L., B.A.J., H.X., Z.A., S.C.W., V.D.M., X.X., P.-Y.S.; Resources, H.X., B.A.J., Z.K., Z.A. X.X.; Data Curation, Y.L., J.L., B.A.J., H.X., Z.K., C.S., S.C.W., V.D.M., X.X., P.-Y.S.; Writing-Original Draft, Y.L., J.L, X.X., P.-Y.S; Writing-Review & Editing, Y.L., J.L., B.A.J., H.X., Z.K., C.S., S.G.W., Z.A., S.C.W., V.D.M., X.X., P.-Y.S.; Supervision, X.X., S.C.W., V.D.M., P.-Y.S.; Funding Acquisition, Z.A., S.C.W., V.D.M., P.-Y.S.

## Competing financial interests

X.X., V.D.M., and P.-Y.S. have filed a patent on the reverse genetic system and reporter SARS-CoV-2. Other authors declare no competing interests.

---

**Extended Data Figure 1.**
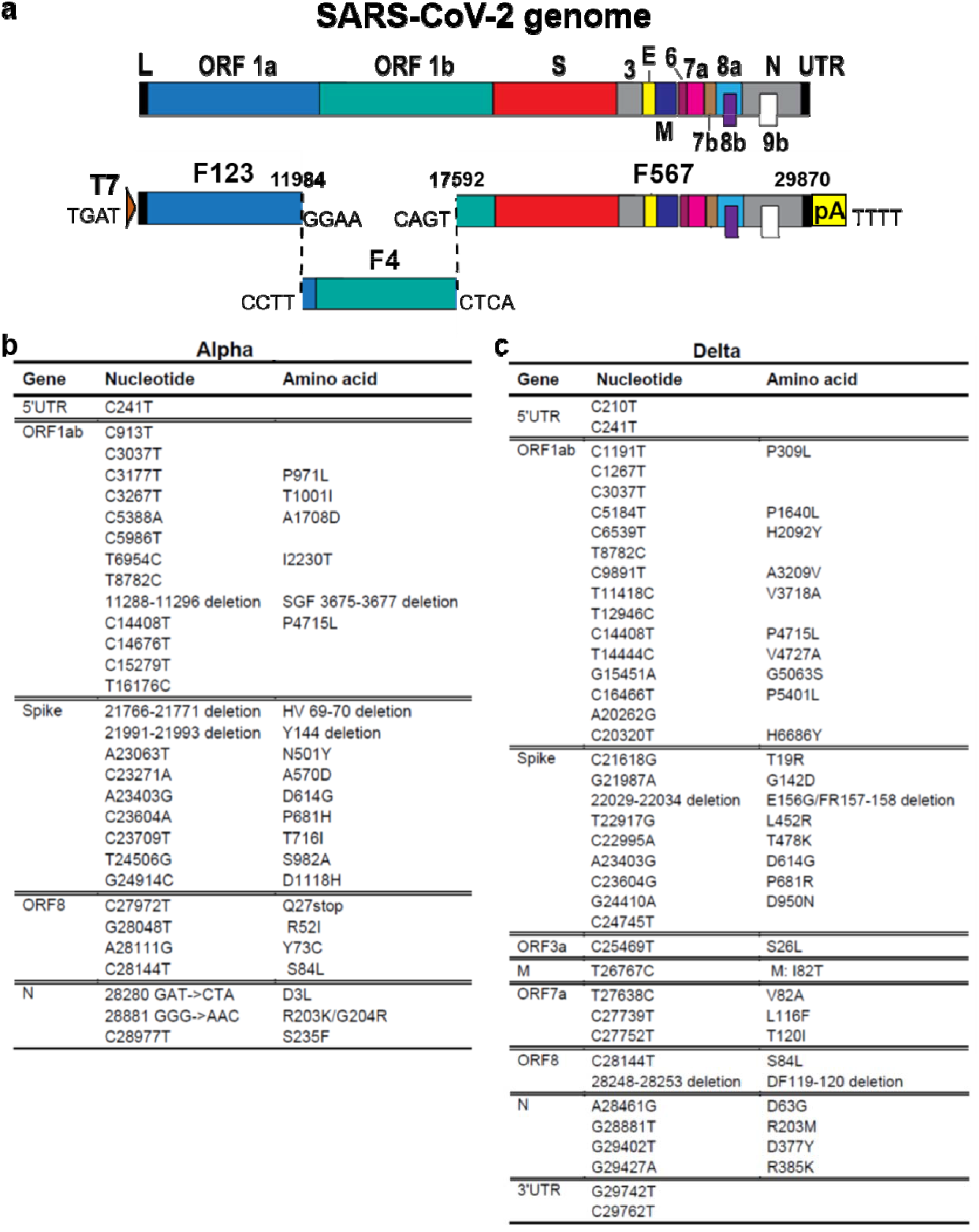
Construction of infectious cDNA clones for Alpha and Delta variants. **a,** Construction of infectious cDNA clones for Alpha and Delta variants. A three-fragment *in vitro* ligation was performed to construct the full-length cDNA clones of Alpha and Delta SARS-CoV-2. The construction method was detailed previously.^12,13^ ORFs, the Open reading frames; L, leader sequence; S, spike gene; E, envelope glycoprotein gene; M, membrane glycoprotein gene; N, nucleocapsid gene; UTR, untranslated region. **b,c,** Mutations from Alpha and Delta variants. The whole genome sequences of Alpha (EPI_ISL_999340) (**b**) and Delta (EPI_ISL_2100646) (**c**) were compared to USA/WA1-2020 strain. Nucleotide and amino acid mutations are presented.

**Extended Data Figure 2.**
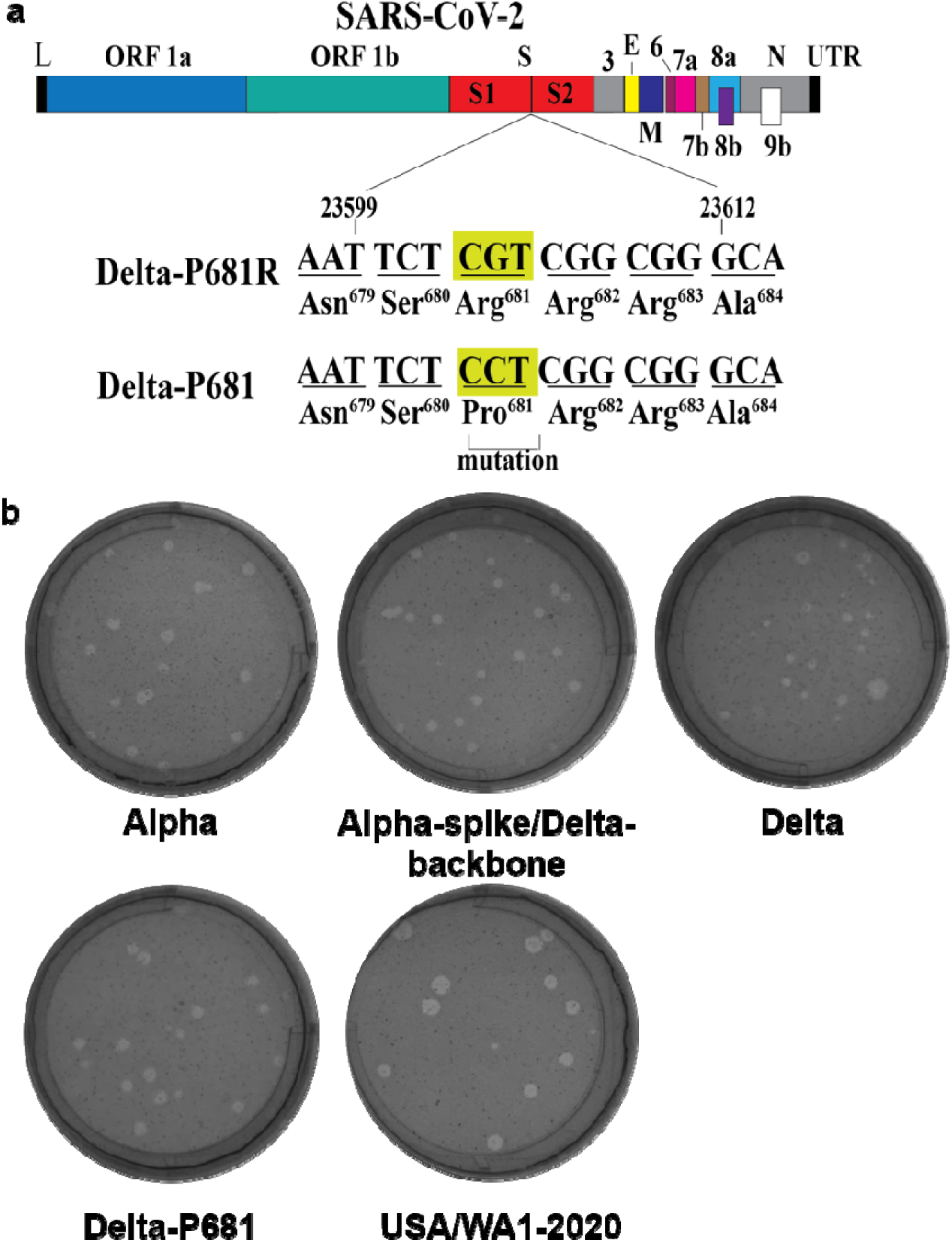
Construction of Delta-P681 SARS-CoV-2 and plaque morphologies of different recombinant SARS-CoV-2s. **a,** Construction of revertant Delta-P681 SARS-CoV-2. Single nucleotide G-to-C substitution was engineered into the Delta variant to construct Delta-P681 SARS-CoV-2. The nucleotide positions of viral genome are annotated. **b,** Plaque morphologies of Alpha, Alpha-spike/Delta-backbone, Delta, Delta-P681, and USA-WA1/2020 viruses. The plaque images were taken on day 2.5 post infection of Vero E6 cells.

**Extended Data Figure 3.**
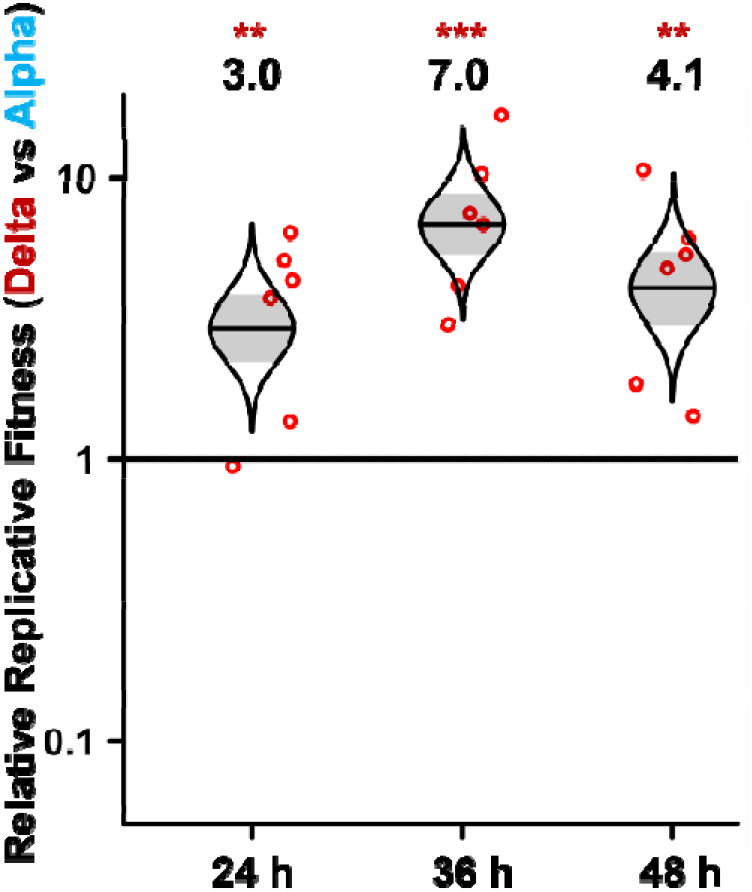
Viral replication competition between Delta and Alpha variants on Calu-3 cells. Recombinant Delta and Alpha SARS-CoV-2s were mixed in equal PFUs to infect Calu-3 cells at a total MOI of 0.1. At 2 h post infection, the cells were washed thrice with DPBS to remove free viruses. Culture medium were sampled for Sanger sequencing at 24 h, 36 h, and 48 h post infection. Red dots represent individual cell cultures (n=6); horizontal lines in each catseye represent the mean; shaded regions represent standard error of the mean; y-axes use a log_10_ scale. Black numbers above each set of values (catseye) indicate the ratios of two viral RNA species. *P* values are calculated for group coefficient using linear regression model. \*\**p*<0.01, \*\*\**p*<0.001.

**Extended Data Figure 4.**
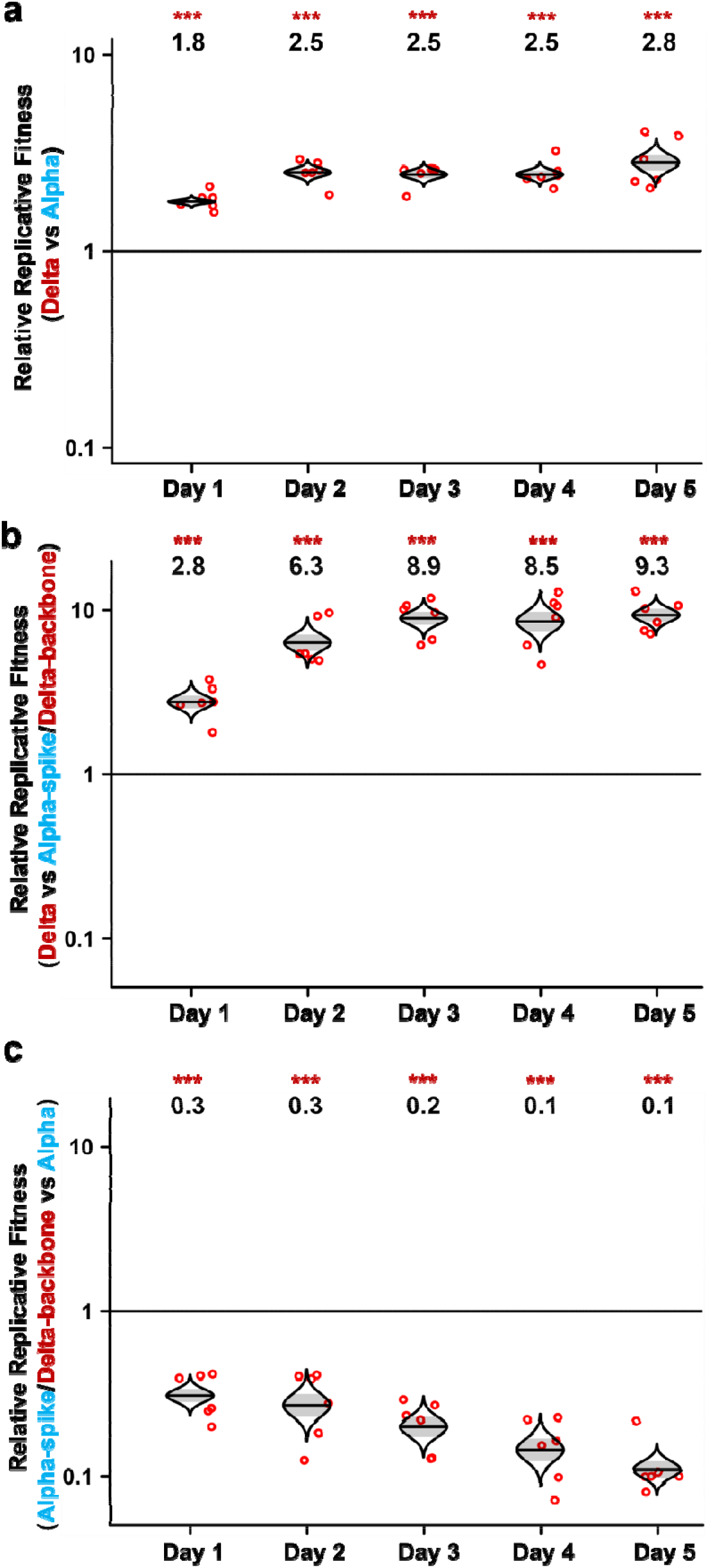
Validation of viral competition results by next generation sequencing. **a-c,** RNA samples from competition assays between Delta and Alpha (**a**), Delta and Alpha-spike/Delta-backbone (**b**), Alpha-spike/Delta-backbone and Alpha (**c**) were initially assessed using Sanger sequencing (**Fig. 1b-d**). The same RNA samples were retested here using next generation sequencing (NGS). Red dots represent individual cell cultures (n=6); the horizontal lines in each catseye represent the mean; shaded regions represent standard error of the mean; y-axes use a log_10_ scale. Black numbers above each set of values (catseye) indicate the relative fitness estimates. *P* values are calculated for group coefficient using linear regression model. \*\*\**p*<0.001.

**Extended Data Figure 5.**
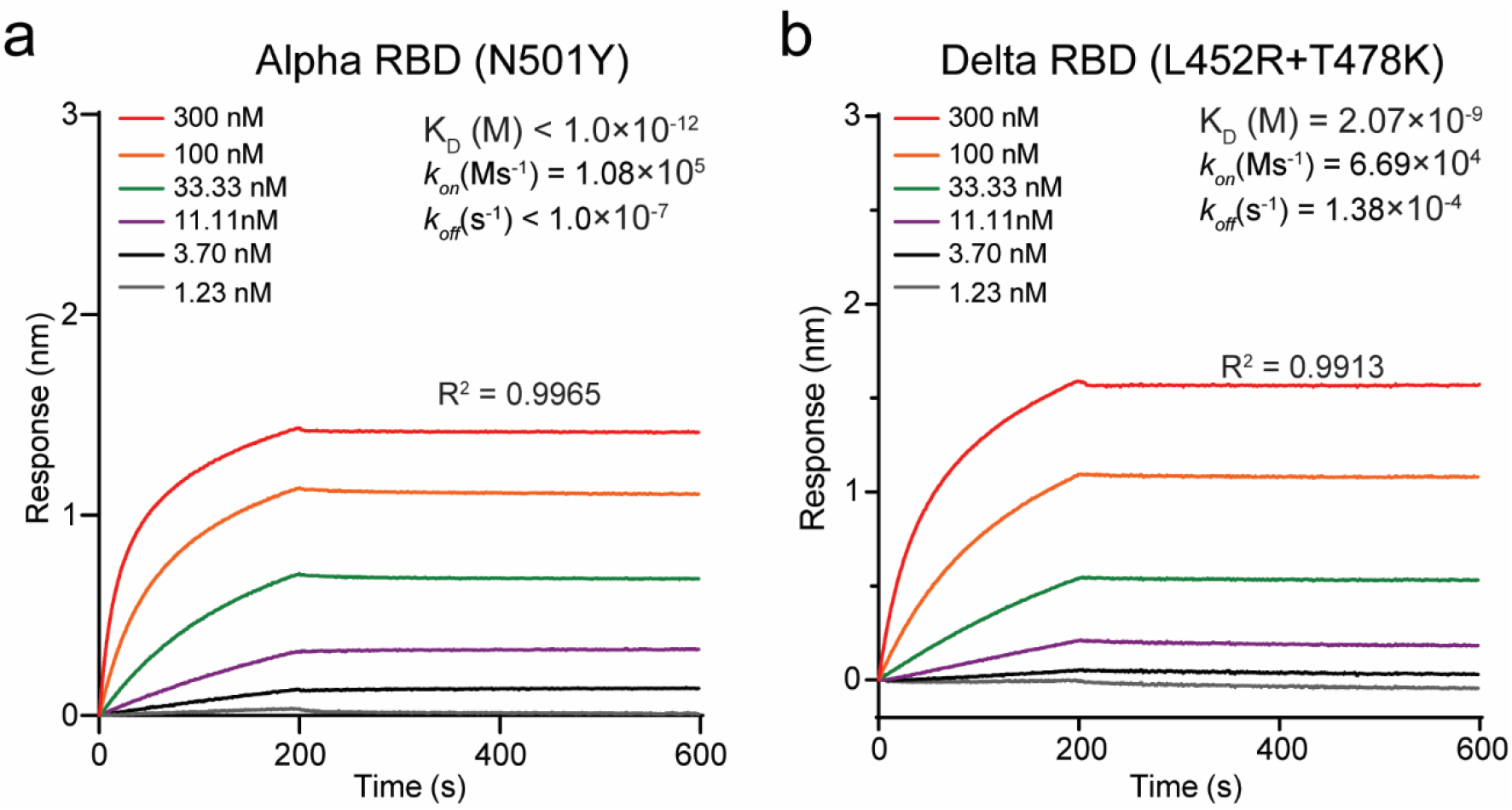
Binding affinities of Alpha and Delta RBDs to human ACE2 receptor. **a-b,** Alpha RBD (**a**) and Delta RBD (**b**) proteins were captured onto protein A biosensors. The biosensors were then dipped into serially diluted human ACE2 protein and buffers to measure the association and dissociation kinetics. The binding affinity-related parameters, including association (K_on_), dissociation (K_off_), and affinity (K_D_) are shown. The affinity of ACE2 to Alpha RBD (N501Y) is below the detection limit and is presented as <1.0×10^−12^. The result for Alpha RBD and ACE2 binding was adopted from our previous study^4^ for comparison.

**Extended Data Table 1.**
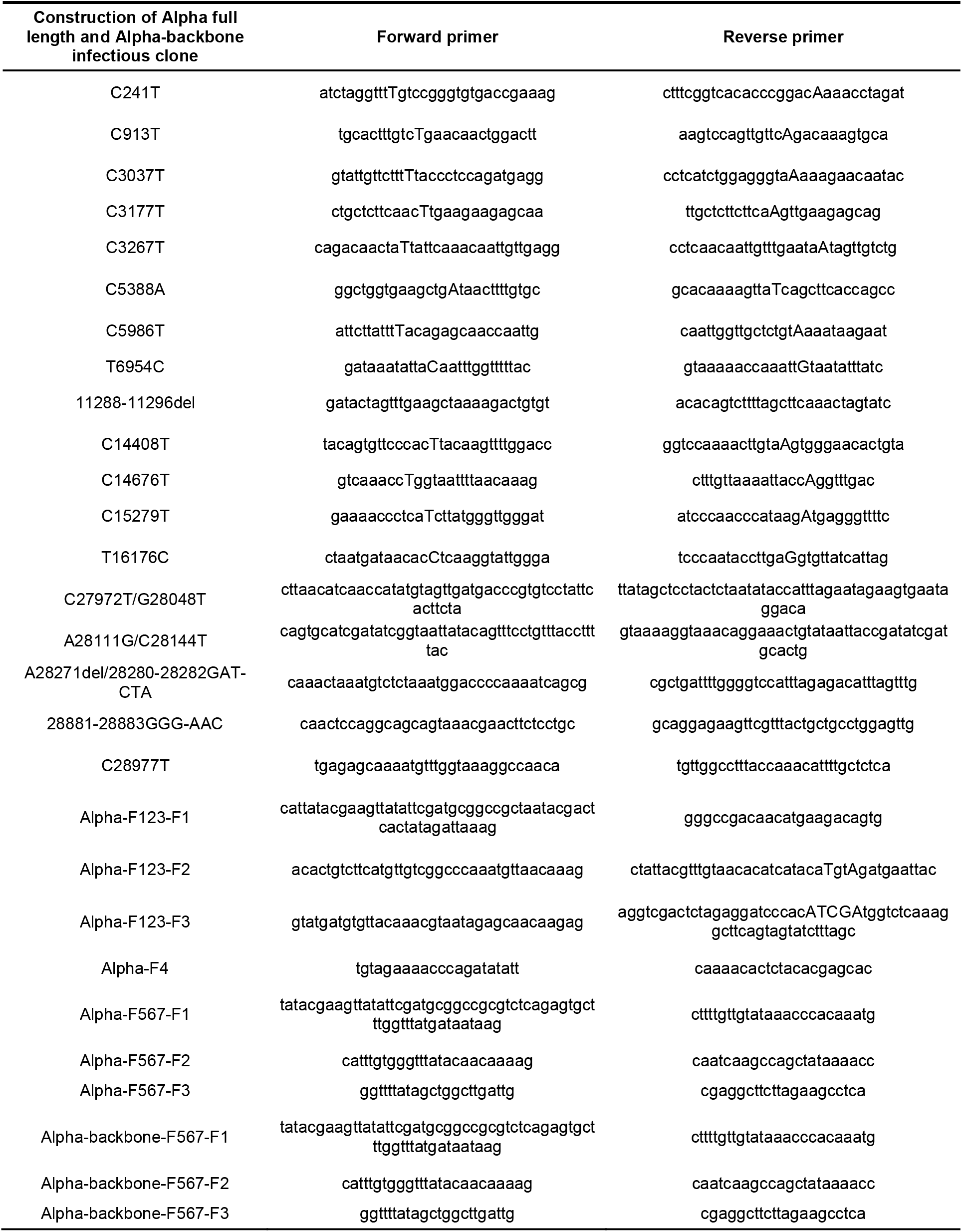
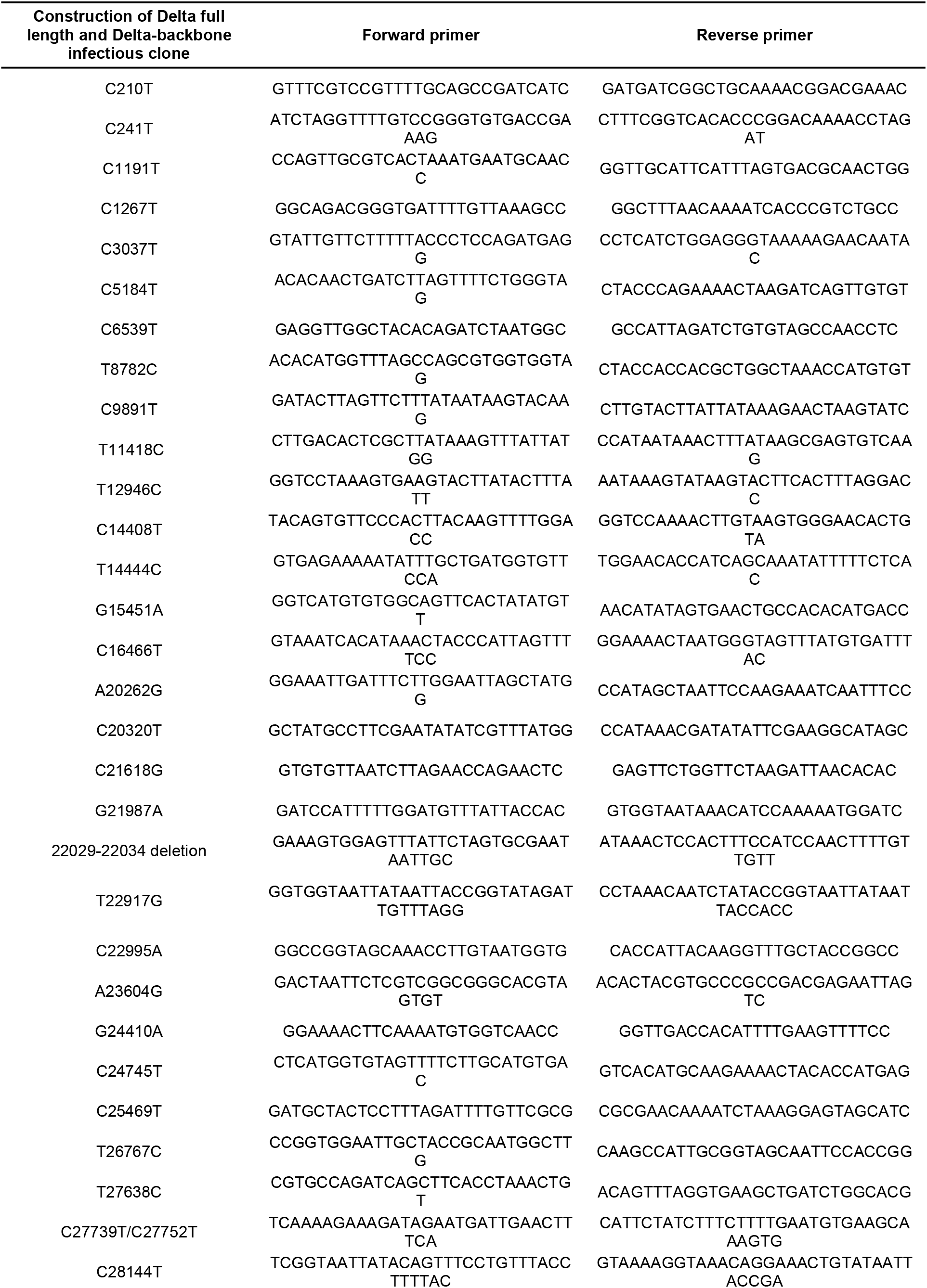
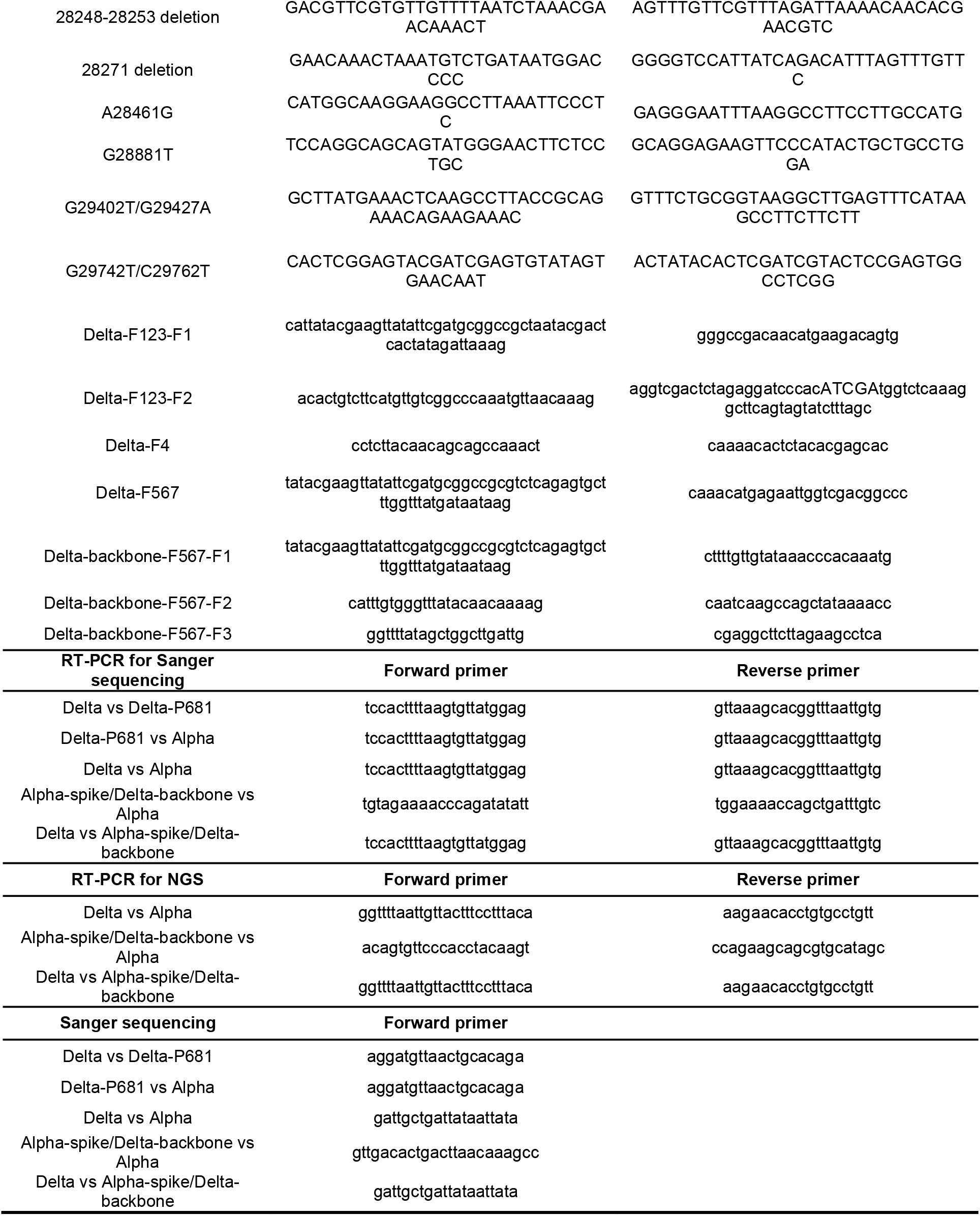
Primer list used for SARS-CoV-2 infectious clones construction, RT-PCR and sequencing.

**Extended Data Table 2.** The input and output viral RNA ratios in competition assays detected by sanger sequencing.

| Groups | HAE |  |  | Calu-3 |  |  |
| --- | --- | --- | --- | --- | --- | --- |
|  | Days | Input ratio | Output ratio | Hours | Input ratio | Output ratio |
| Delta vs Alpha | Day 1 | 1.24 | 2.10 | 24h | 1.53 | 4.56 |
|  | Day 2 | 1.24 | 3.24 | 36h | 1.53 | 10.65 |
|  | Day 3 | 1.24 | 3.17 | 48h | 1.53 | 6.21 |
|  | Day 4 | 1.24 | 3.51 |  |  |  |
|  | Day 5 | 1.24 | 3.78 |  |  |  |
| Delta vs Alpha-spike/Delta-backbone | Day 1 | 1.07 | 2.97 |  |  |  |
|  | Day 2 | 1.07 | 6.93 |  |  |  |
|  | Day 3 | 1.07 | 10.60 |  |  |  |
|  | Day 4 | 1.07 | 9.64 |  |  |  |
|  | Day 5 | 1.07 | 10.53 |  |  |  |
| Alpha-spike/Delta-backbone vs Alpha | Day 1 | 1.45 | 0.59 |  |  |  |
|  | Day 2 | 1.45 | 0.46 |  |  |  |
|  | Day 3 | 1.45 | 0.28 |  |  |  |
|  | Day 4 | 1.45 | 0.22 |  |  |  |
|  | Day 5 | 1.45 | 0.15 |  |  |  |
| Delta vs Delta-P681 | Day 1 | 0.95 | 2.05 |  |  |  |
|  | Day 2 | 0.95 | 2.20 |  |  |  |
|  | Day 3 | 0.95 | 2.38 |  |  |  |
|  | Day 4 | 0.95 | 2.77 |  |  |  |
|  | Day 5 | 0.95 | 3.25 |  |  |  |
| Delta-P681 vs Alpha | Day 1 | 1.51 | 1.33 |  |  |  |
|  | Day 2 | 1.51 | 1.13 |  |  |  |
|  | Day 3 | 1.51 | 1.21 |  |  |  |
|  | Day 4 | 1.51 | 0.97 |  |  |  |
|  | Day 5 | 1.51 | 0.85 |  |  |  |

